# Revealing determinants of translation efficiency via whole-gene codon randomisation and machine learning

**DOI:** 10.1101/2022.04.05.486962

**Authors:** Thijs Nieuwkoop, Barbara Terlouw, Dick de Ridder, John van der Oost, Nico J. Claassens

## Abstract

Codon usage refers to the occurrence of synonymous codons in protein-coding genes. It is known for decades that codon usage contributes to translation efficiency and hence to protein production levels. However, its role in protein synthesis is still only partly understood. This lack of understanding hampers the design of synthetic genes for efficient protein production. In this study, we developed a method to generate a large, synonymous codon library of the gene encoding the red fluorescent protein (RFP). After expression in *Escherichia coli*, 1459 clones of this library were selected of which we measured protein production levels and determined the full coding sequences. Using different machine learning approaches, this data was used to reveal correlations between codon usage and protein production. Interestingly, protein production levels can be relatively accurately predicted (Pearson correlation of 0.762) by a Random Forest model, which only relies on the sequence information for the first 8 codons. This study clearly demonstrated the key role of codons at the start of the coding sequence. As such, it provides not only important fundamental insights on the influence of codon usage on protein production but also relevant clues on optimising the design of efficiently translated synthetic genes.

## INTRODUCTION

Due to degeneracy in the genetic code, a protein with a single amino acid sequence can be encoded by an extremely large number of different coding sequences (CDS). While different synonymous codons do not alter the amino acid sequence, they are known to influence translation efficiency and in some cases even protein folding properties (1–5). However, many questions about the roles of codons and their often subtle and intertwined effects are still unanswered. Understanding codon usage is key to grasping one of the fundamental processes of life: the translation of mRNA into proteins. In addition, precise control over translation efficiency is highly desirable in both biotechnology and synthetic biology to make the process of protein production and cell engineering more predictable.

Since the early days of DNA sequencing it was observed that, depending on the organism, specific codons are overrepresented (6). This led to the hypothesis that frequently occurring codons could be translated more efficiently, e.g., due to a higher abundance of corresponding tRNAs. It also led to the so-called Codon Adaptation Index (CAI) (6), which is defined as the geometric mean of the relative codon usage in a specific coding sequence (based on the average codon usage in the genome or a subset of highly expressed genes). In other words, a CDS with a high CAI primarily uses frequent codons, while a CDS with a low CAI contains more rare codons. However, the hypothesis that a high CAI is related to high protein production has been disputed in several studies in recent years (2, 4, 7). Especially the hallmark study by Kudla *et al.*, in the bacterium *Escherichia coli*, revealed that the CAI does not seem a major determinant of high protein production. In this study, a set of 154 codon variants of the Green Fluorescent Protein (GFP) was generated. The authors could not correlate the CAI to the protein expression levels. However, the predicted folding energy of the mRNA around the start codon did correlate with protein production. They hypothesised that the expression efficiency in *E. coli* is mostly influenced by the availability of the ribosome binding site (RBS) for translation initiation. Several studies have since followed up on this and agreed on a key role of mRNA secondary structures (4, 8–10) around the RBS and start codon. The codon usage at the 5’ CDS has also been hypothesised to be involved in a so-called codon ramp, as these regions generally contain more rare codons (11, 12). This ramp would result in a slow initial translation elongation speed, reducing the risk of detrimental ribosomal collisions along the length of the CDS. Finally, codon usage has also been associated with mRNA stability (1, 3, 13). Especially in eukaryotes, slow-moving ribosomes can initiate RNA decay, thereby linking translation elongation efficiency to mRNA decay (14).

Despite a range of studies in this field, however, there are still many open questions about the complex role of codon usage on expression in different organisms. Consequently, reliable models for predicting protein production based on codon usage are unavailable. Many of the algorithms used for ‘codon optimisation’ are based on the CAI score or variations thereof, which in practice often fail to give optimal results (15).

To further contribute to the understanding and predictability of protein production based on codon usage, we decided to generate a large gene-wide synonymous codon library of the gene encoding the monomeric Red Fluorescent Protein (mRFP), and express these in *E. coli*. This reporter protein has hardly been used in studies that focus on codon usage, as opposed to GFP. We chose a different reporter protein to see if a different gene candidate would lead to new findings on the determinants of codon usage. To improve our fundamental understanding of the impact of codon usage, and in an attempt to improve the predictability of optimal codon usage for protein production, we decided to test different machine learning approaches. Very recently, some studies have successfully utilised machine learning methods to predict gene expression based on randomised sequence libraries for non-coding gene regions, such as promoters and 5’ untranslated regions (5’ UTR) in *E. coli* and *Saccharomyces cerevisiae* (16, 17).

In this study, we constructed *mRFP* gene libraries for which the codon usage throughout almost the whole gene was fully randomised. To this end, we developed an assembly approach based on type IIS restriction and ligation. After library assembly, high-quality curated protein production levels were measured for 1459 individual clones, and their specific coding sequences were accurately determined. We then used these pairs of CDSs and corresponding expression values as training data for our machine learning algorithm MEW (mRNA Expression Wizard), to establish an algorithm that can predict the expression from the CDS. Remarkably, we show that only a window covering codons 2-8 is required to accurately predict mRFP production, based on sequence information only. This further strengthens the conclusions from previous studies and underlines that future studies aiming to understand or engineer protein production should focus on the codon usage of the 5’ start of the coding sequence.

## MATERIALS AND METHODS

### mRFP codon randomisation

The amino acid sequence of the monomeric Red Fluorescent Protein (mRFP) was used to generate three degenerate DNA sequences representing our libraries (CAI_L_, CAI_M_ and CAI_H_). Each degenerate sequence was then split into blocks of roughly equal sizes (80-90 nucleotides) in such a way that each block has a unique 4-base pair overlap with neighbouring blocks. Overhangs were selected from a set that is optimised for high ligation fidelity (18). To create each required overhang, we attempted to fix degenerate codons in such a way that the separate blocks were roughly equal in size, and that loss of degeneracy stayed limited. For example, fixing the degenerate sequence ARAT to AAAT would result in the loss of 1 codon possibility, while fixing the degenerate sequence YGCN to CGCC would result in the loss of 7 codon possibilities. 5’ and 3’ flanking sequences containing recognition sites for the Type II restriction enzyme BsaI-HF^®^v2 (NEB, R3733) were added to each IUPAC DNA block, to generate the unique single-stranded overhang after digestion. The 5’ end of the first block and the 3’ end of the last block contained SapI (NEB, R0569) recognition sites instead. Each block was ordered as a DNA oligo (Ultramer^®^ DNA Oligonucleotides, IDT) and using a strand-displacing Taq polymerase (NEB, M0482), the ssDNA was converted to double-stranded DNA via PCR. PCR reactions containing the dsDNA block were cleaned and concentrated to 20 μl mQ using the DNA Clean & Concentrator^™^-5 kit (Zymo, D4004). 4 μl Gel Loading Dye, Purple (6x) (NEB, B7024) was added to each block and they were loaded on a 1% agarose gel and ran for 30 minutes at 100 volts. The dsDNA blocks were excised from the gel and purified to 20 μl mQ using the Zymoclean^™^ Gel DNA Recovery Kit (Zymo, D4002). 5 μl of the dsDNA was used to quantify the DNA concentration with the Qubit assay (Invitrogen, Q32853) according to the manufacture’s protocol.

The dsDNA blocks were mixed in an equal molar ratio to a total volume of 41 μl, with 5 μl T4 Ligase Buffer (NEB, B0202), 400 units T4 Ligase (NEB, M0202) and 60 units BsaI-HF^®^v2 (NEB, R3733). Assembly reaction was done overnight at 37 °C for 18 hours, followed by 5 minutes at 60 °C and a holding step at 12 °C. The assembly is cleaned and concentrated to 15 μl mQ using the DNA Clean & Concentrator^™^-5 kit (Zymo, D4004). 3 μl Gel Loading Dye, Purple (6x) (NEB, B7024) was added and the assembly mixture was loaded on a 1% agarose gel and ran for 40 minutes at 100 volts. The fulllength assembled product was excised from the gel and purified to 44 μl mQ using the Zymoclean^™^ Gel DNA Recovery Kit by Zymo (D4002). 10 units of SapI (NEB, R0569) was added with 5 μl CutSmart Buffer (NEB, B7204) and digested for 2 hours at 37 °C. The digested codon random mRFP with singlestranded overhangs was cleaned and concentrated to 15 μl mQ using the DNA Clean & Concentrator^™^-5 kit by Zymo (D4004). The complete 15 μl containing the codon random mRFP library was used in a ligation reaction to generate the plasmid library.

### Plasmid preparation and library generation

The pFAB3909 plasmid (19) (Addgene #47812) with a P15A origin, kanamycin resistance gene and bicistronic design element was modified to be able to accept the codon randomised mRFP library and include a constitutively expressed GFPuv gene. The relatively weak bla promoter was used to drive the mRFP expression, keeping the total protein yield relatively low to prevent expression saturation for high producing mRFP codon variants. A strong terminator was used for efficient transcription termination and to enhance mRNA stability. The open reading frame was replaced by SapI recognition sites to generate the sticky overhangs that accept the mRFP library and a large part of nonsense DNA was inserted between the SapI sites to separate the double SapI digested plasmid from linear product. A GFPuv gene, driven by the P4 promoter, was added to the plasmid as an internal standard for gene expression. Expression of GFPuv is weak as to not interfere with the mRFP expression efficiency but strong enough for detection with flow cytometry.

About 3 μg plasmid was digested with 20 units SapI (NEB, R0569) and dephosphorylated with 3 units rSAP (NEB, M0371) with 6 μl CutSmart Buffer (NEB, B7204) in a total volume of 60 μl for 3 hours at 37 °C, followed by an inactivation step at 65 °C for 20 minutes. The linear plasmid was excised from the gel and purified to 30 μl mQ using the Zymoclean^™^ Gel DNA Recovery Kit by Zymo (D4002). The codon random mRFP library (15 μl) was ligated into 30 ng linear plasmid with 400 units of T4 ligase (NEB, M0202) and 2 μl T4 Ligase Buffer (NEB, B0202) in a total volume of 30 μl for 18 hours at 16 °C. The ligation mixture was cleaned and concentrated to 10 μl mQ using the DNA Clean & Concentrator^™^-5 kit by Zymo (D4004). 1 μl of the codon randomised mRFP library was transformed into electrocompetent DH10B cells (20 μl competent cells, 2mm cuvette, Voltage: 2500V, Resistor: 200 Ω, Capacitor 25 μF, BTX^®^ ECM630). Cells were recovered in 1 ml NEB^®^ 10-beta/Stable Outgrowth Medium (NEB, B9035) at 37 °C for 1 hour. The cells were transferred to a 50 ml tube and 9 ml LB (10 g/l Peptone (OXOID, LP0037), 10 g/l NaCl (ACROS, 207790010) and 5 g/l Yeast Extract (BD, 211929)) were added with 50 μg/l kanamycin (ACROS, 450810500) and incubated for 18 hours at 37 °C.

### Expression range enrichment and selection

A FACS (Sony, SH800S Cell Sorter; GFPuv excitation at 488 nm, emission at 525/50 nm; mRFP excitation at 561 nm, emission at 617/30 nm) was used to sort 50,000 cells of the overnight cell culture into 3 groups based on expression. The left and right tail of the normal distribution and a part of the middle peak was sorted to create 3 groups of low, medium and high expression. The 3 cell groups were put on individual agar plates (10 g/l Peptone (OXOID, LP0037), 10 g/l NaCl (ACROS, 207790010), 5 g/l Yeast Extract (BD, 211929), 15 g/l Agar (OXOID, LP0011), 50 mg/l kanamycin (ACROS, 450810500)) and grown overnight at 37 °C. From these plates, individual colonies were picked and grown in 2 ml 96 well plates with 200 μl LB with kanamycin (10 g/l Peptone (OXOID, LP0037), 10 g/l NaCl (ACROS, 207790010), 5 g/l Yeast Extract (BD, 211929), 15 g/l Agar (OXOID, LP0011), 50 mg/l kanamycin (ACROS, 450810500)) for 18 hours at 37 °C.

### Measurements and sequencing

The cell cultures were diluted 100x in PBS (8 g/l NaCl (ACROS, 207790010), 200 mg/l KCl (ACROS, 196770010), 144 mg/l Na2HPO4 (ACROS, 12499010), 240 mg/L KH2PO4 (ACROS, 447670010)). mRFP expression was measured using a flow cytometer (Thermo, Attune NxT Flow Cytometer; GFPuv excitation at 405 nm, emission at 512/25 nm; mRFP excitation at 561 nm, emission at 620/15 nm; stop option 200,000 single cells). A gate was used to exclude GFPuv outliers (±10% of the total population) aiming to reduce unrelated biological variance as GFPuv expression levels are expected to stay constant. From the overnight cultures, 1 μl of cells were used in a PCR reaction to obtain the gene for Sanger sequencing using Q5 (NEB, M0492). The PCR reaction was sent to Macrogen Europe B.V. for sample clean-up and Sanger sequencing.

All cell cultures were also measured using a microplate reader (BioTek, Synergy Mx). 50 μl overnight cell cultures were diluted in 50 μl PBS (8 g/l NaCl (ACROS, 207790010), 200 mg/l KCl (ACROS, 196770010), 144 mg/l Na2HPO4 (ACROS, 12499010), 240 mg/L KH2PO4 (ACROS, 447670010)). The plates were incubated at room temperature for 1 hour before measuring (cell density measured at 600 nm; GFPuv excitation at 395/9 nm, emission at 508/9 nm; mRFP excitation at 584/9, emission at 607/9 nm). The microplate reader fluorescent readings were normalised with the OD600 for both the GFPuv and mRFP readings.

### Data validation

Before using the measurements in our machine learning approach, we set a few criteria that the data had to meet in order to exclude artefacts in our dataset. First, the sequencing data should be of sufficient quality and the encoded amino acid sequence should be correct. The raw sequence data was validated by extracting the open reading frame sequence using in-house scripts. All bases in the open reading frame needed a Phred quality score > 20 (a base call accuracy of at least 99%) and the translated sequence should match the mRFP amino acid sequence in order to pass. Second, any double populations or clear changes in cell morphology or culture density were excluded. Double populations were apparent in the flow cytometry data (see figure S2A) but were already automatically excluded due to the sequencing quality criteria as a double population will result in poor Sanger sequencing data. However, very rarely, we observed an unexplained shift in cell morphology as increased forward and side scatter values were obtained during flow cytometry (figure S2B). Finally, rarely, a difference was observed in fluorescence between our flow cytometry measurements and microplate reader measurements. Based on an arbitrary threshold of 25% deviation from the average relationship between the two measurement methods we excluded these deviating cell cultures (see Methods and Figure S3). All exclusions were made prior to our analysis in an attempt to generate high-quality data to feed the machine learning algorithm. For the remaining data points, we assessed dataset-wide biases and correlations, such as assembly bias and the correlation between expression level and GC content to ensure the dataset as a whole was appropriate for machine learning. All raw data and validated data are available in Supplementary Data S1 and S2.

### Building machine learning regressors

To assess if mRNA expression levels could be predicted from sequence, we employed two different machine learning approaches: Random Forest Regressor (RFR) and LASSO. We implemented RF and LASSO using the scikit-learn package (v0.23.0, ref) in python (v3.7.6), with the sklearn.ensemble.RandomForestRegressor and sklearn.linear_model.Lasso modules respectively. For RF, default settings were used, while for LASSO, various values for alpha were assessed (0.05, 0.1, 0.5, 1.0, 2.0, 5.0, 10.0, 20.0, 50.0, and 100.0 for all regressors, in addition to 500.0 and 1000.0 for regressors trained on full-length sequences; max iterations = 10,000). An alpha of 100.0 gave rise to the best predictions for full-length sequences (Figure S4); for windows, different alphas performed better for different featurisations, but differences were minimal. Separate regressors were constructed for fulllength featurised mRNA sequences and for featurised sliding windows of 10, 20, 30, or 40 bases. Prior to training, an independent test set comprising 10% of the data was set aside. Regressor accuracies were then evaluated on the remaining training data through 10-fold cross-validation, where 90% of the training data were used to predict the translation efficiency of the other 10%. This was done for each 10% of the data, such that we obtained a predicted translation efficiency, measured with flow cytometry, for each data point. From these predictions, Pearson and Spearman correlations were computed for actual flow vs predicted flow and used as measures for model accuracy. Feature importances were extracted from all ten regressors built in cross-validation, averaged, and plotted and visualised with matplotlib (v3.2.1). Finally, for regressors trained on the full-length sequences and the best-performing sequence windows (Figure S5, Figure S6), new regressors were trained using all training data, and model accuracies were re-evaluated on the independent test-set. Code and regressors are made available at https://github.com/BTheDragonMaster/mew.

## RESULTS AND DISCUSSION

### Type IIS assembly method allows for generation of codon randomised libraries at different CAI levels

To randomise synonymous codon usage throughout the whole *mRFP* CDS, we developed a new randomisation-assembly method based on type IIS restriction and ligation. We used this approach to generate three codon randomised *mRFP* libraries with either fully randomised codon usage or with a focus on more frequent or rare codons. The first CAI library (“medium”, CAI_M_) is fully randomised and uses an equal distribution of all synonymous codons for each amino acid. Only for the amino acids arginine, leucine and serine, as well as the stop codons, the full codon space could not be covered due to the sequence limitations in degenerate oligos. For each of the three amino acids 4 of the 6 possible codons were included, while the stop codon was kept constant at TGA. The theoretically maximum number of CDSs coding for the mRFP protein is 3.19 × 10^104^. By limiting the aforementioned amino acids, the stop codon and a few codons needed for assembly purposes the library still contains at most 3.68 × 10^93^ variants. Obviously, this is an astronomically large number and generated libraries and experimental efforts can only cover a very small fraction of this diversity.

The randomised design approach results in a uniform codon bias distribution across the gene, with an overall medium CAI of 0.67 (Figure S1). To see if codon randomisation with an overrepresentation of frequent or rare codons affects translation differently, we generated two additional libraries. By restricting the allowed relative adaptiveness (the usage ratio of a codon to that of the most abundant synonymous codon), we generated libraries that use more frequent or rare codons. The library limited to rare codons used only synonymous codons with a relative adaptiveness <0.60, or the lowest relative adaptiveness in the case the synonymous codons are used in a close to equal ratio. This rare codon library (CAI_L_) had an overall CAI of 0.41 (Figure S1). The library using frequent codons (CAI_H_) used only synonymous codons with a relative adaptiveness >0.50, resulting in a library with an overall CAI of 0.83 (Figure S1). The theoretical numbers of possible sequences for the libraries are listed in Table S1.

To create the three libraries, we divided the complete degenerate CDS into eight blocks of ~85 bases, which can be assembled with unique overhangs between each adjacent part. In order to generate complementary four base pair overhangs between the parts, some codons with multiple synonymous options needed to be fixed to a single codon. The eight DNA parts were ordered as single-stranded oligos with additional Type IIS restriction sites flanking the blocks. The oligos were converted to doublestranded DNA using PCR and consequently assembled using Type IIS restriction and ligation (Figure 1A, B). Only a small fraction of the total DNA parts assembled into the full product of 707 bp (Figure 1B, indicated with the “*mRFP*” label). Seven intermediate products were observed that did not further assemble into the full gene. The assembly limitations could originate from synthesis errors in the initial oligos, preventing Type IIS restriction or resulting in incorrect overhangs. The band corresponding to the fully assembled product was purified and ligated into an expression vector (Figure 1C).

**Figure 1.**
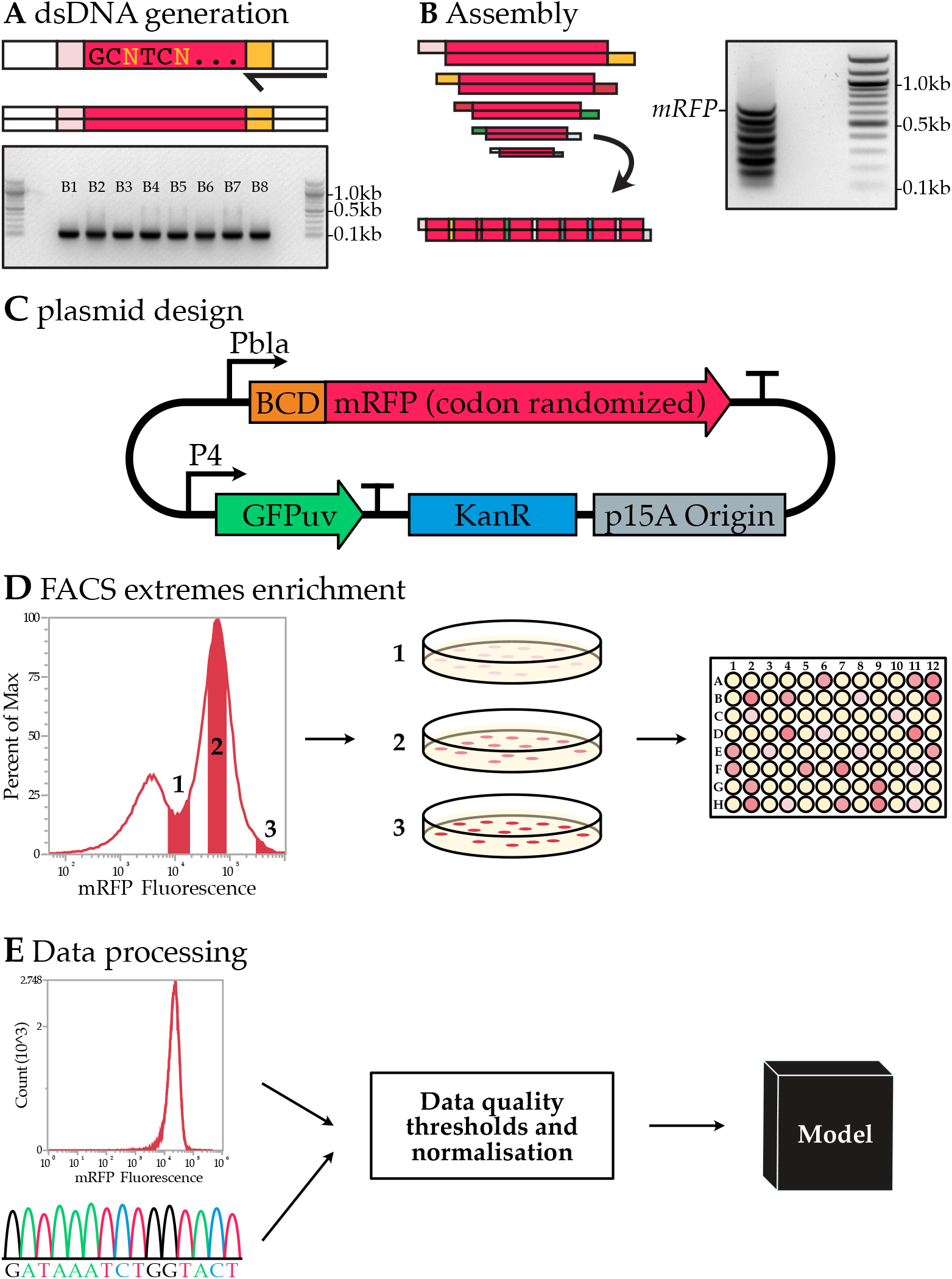
Codon randomised library generation and analysis. (**A**) PCR is used to generate dsDNA from oligos and an electrophoresis gel yields the eight dsDNA blocks used to build codon randomised mRFP. (**B**) The assembly reaction and the electrophoresis gel result of the assembly. The complete assembly of all eight blocks is indicated with the *mRFP* tag. The seven bands below are intermediate products. (**C**) The expression vector used to express the codon random mRFP. (**D**) FACS enrichment for a wide expression range within the library is used to obtain a higher representation of the high and low expressing codon variants. (**E**) Flow cytometry analysis of cultures and Sanger sequencing data are QA passed and used in machine learning models.

The expression vector contains a native, relatively weak, beta-lactamase promoter (Pbla). A weak promoter will pose less burden on the transcriptional and/or translational processes, reducing the risk of reaching an upper limit in the protein production process and thus preserving the full expression range as determined by the coding sequence. For the 5’ UTR, a medium-strength bicistronic design (BCD) was chosen based on the work of Mutalik et al. (20). This BCD element (BCD5) was previously reported to reduce the influence of mRNA secondary structures on expression. Including this element allows us to study the more nuanced features associated with codon usage and rules out strong effects of mRNA structure formation with the constant 5’ UTR.

### Expression of libraries in *E. coli* results in wide expression range and allows for high-quality data collection

The library containing codon randomised *mRFP* was transformed into *E. coli* DH10B. A single transformation of the libraries in *E. coli* yielded between 150.000 and 320.000 colonies. After 18-hour cultivation on liquid, roughly 70% of the cells gave a detectable level of red fluorescence (measured using flow cytometry, Figure 2). The remaining 30%, for which no or very little fluorescence was measured, was later confirmed via sequencing to mainly comprise constructs that had a frameshift in the ORF. This is not unexpected, as some blocks are likely missing one or multiple nucleotides since the coupling efficiency of oligos is not 100%. These errors eventually lead to frameshifts and thus protein truncations or mutations.

**Figure 2.**
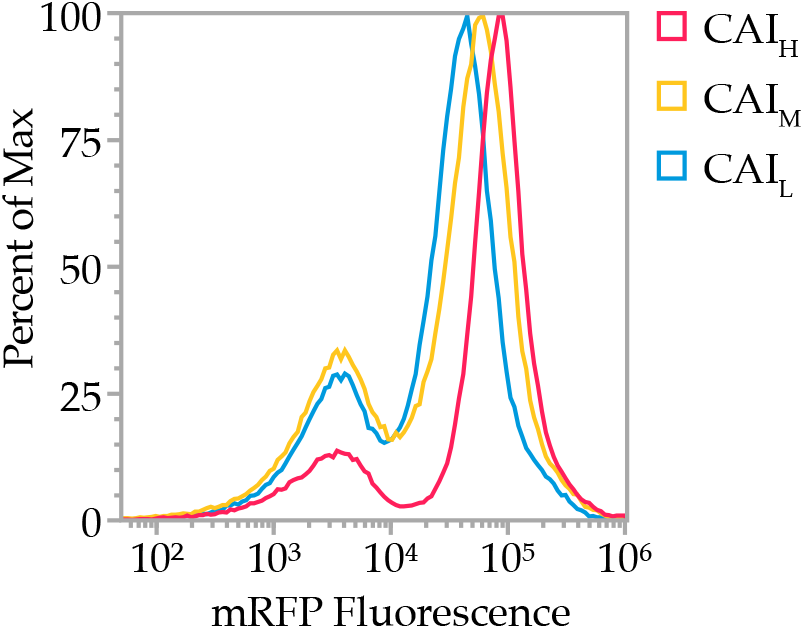
Normalised flow cytometry overlay of the mRFP fluorescence signal from the CAI_L_, CAI_M_ and CAI_H_ libraries. The left peak is part of the population showing no fluorescence, mainly due to assembly errors in the CDS. The right peak shows the mRFP expression of each library. The average expression of the CAI_L_, CAI_M_ and CAI_H_ libraries is in increasing order, but high expressing variants are found in all libraries (right tail). The ratio between the left and right peaks shows the fidelity of the library as the left peak consists of autofluorescence of non-expressing or non-functional variants.

The flow cytometric evaluation of the three library populations showed that the average expression of the CAI_L_, CAI_M_ and CAI_H_ libraries was in increasing order (Figure 2). This suggests that an overall higher CAI leads to higher expressing constructs on average. However, for all three libraries, expression could be observed at the highest end of the expression spectrum, suggesting that a high CAI is not the leading factor for high protein production.

To obtain high-quality expression and sequence data for the downstream machine learning analyses, we decided to study expression levels and related sequences of individual clones. We favoured this method over previously used FlowSeq methods, which perform sequencing analysis on large mixes of clones obtained during fluorescence activated cell sorting (FACS). FlowSeq typically employs shortread sequencing (Illumina), which will not cover the full CDS length in a single read and due to the whole-gene codon degeneracy in this study, it would be difficult to assemble reads into contigs. Alternative long-read single-molecule methods (e.g., PacBio) would offer a solution, but it was questionable whether a sufficiently high coverage could be achieved to reach meaningful conclusions. FlowSeq has another limitation, as the fluorescence level detected from cells with the same genotype already can cover a relatively wide range (21). This increases the likelihood that individual cells are binned incorrectly and that the resulting dataset is too noisy to be analysed meaningfully through statistical analyses and machine learning. This can potentially be solved by sequencing with a very high coverage; however, this is hard to achieve for high-quality long reads as mentioned above. We chose to select a limited number of individual clones for which mean expression values can be accurately determined, as well as their full gene sequences using Sanger sequencing.

To allow the selected clones to cover a wide range of low, medium and high expressing constructs, and exclude non-expressing (e.g., frameshifted constructs), a preselection of three groups was first performed using FACS (Figure 1D). After sorting in these bins, we picked colonies from the three different expression-level groups for each CAI library. These clones were all inoculated in liquid culture (for a total of 480, 1440 and 480 individual cultures for CAI_L_, CAI_M_ and CAI_H_ respectively). The fluorescence of these cell cultures was measured using both flow cytometry and a microplate reader. The mRFP expression was normalised with the constant constitutively expressed GFP (Figure 1C). The *mRFP* coding sequence (and untranslated regions) was amplified using colony PCR and amplicons were analysed by Sanger sequencing. Next, the data was evaluated to exclude low-quality sequencing reads, amino acid mutations, mixed populations (Figure S2A), and rare deviations in cell morphology (notable increase in FSC and SSC as observed in the flow cytometer, Figure S2B) or culture density (deviation between measurements with flow cytometry and microplate reader, Figure S3). The exclusion criteria are further described in the Materials and Methods – Data validation section. This left us with 1459 high-quality data points that we could use in our machine learning approach (Figure 1E).

### Different machine learning approaches can predict mRPF production levels

To identify the determinants of protein production levels and to assess if the expression levels could be predicted from gene sequence, we employed two different machine learning approaches: Random Forest Regressor (RFR) and LASSO (Least Absolute Shrinkage and Selection Operator). For this purpose, we developed MEW: the mRNA Expression Wizard, which can train and test a variety of machine learning models to predict the protein expression level from mRNA sequence, using different types of featurisations. These featurisations include methods that focus on the base pair composition of the coding sequence, to observe the effects of factors like translation elongation efficiency, and vectorisations that reflect the probability that a base is paired in the context of an mRNA secondary structure.

Our rationale to use both LASSO and RFR is that due to their stepwise decision making, RFRs can model non-linear interdependencies between bases, while LASSO is better suited to straight-forward linear regression and feature selection. Importantly, for each regressor we trained we extracted the feature importances as this could help to identify determinants of translation efficiency. We trained separate regressors for both full-length featurised mRNA sequences and for sliding windows of varying sizes along the entirety of the mRNA, to assess if certain windows are more predictive of translation efficiency than others.

As performance of machine learning algorithms depends greatly on their input data, featurising our mRFP data in a way that captures most information was key. We used three featurisation methods: one based on one-hot encoding of base identity, another based on predicted pairing probabilities of bases in mRNA secondary structure calculated with ViennaRNA (22), and a third which combines these two. While one-hot encoding should in theory also be able to capture base pairing probability, we decided to also use mRNA secondary structure featurisation as the resulting features are easier to interpret. We will call the three types of featurisations BPP (base pairing probability), one-hot, and BPP + one-hot respectively.

We trained and validated the RFR and LASSO regressors with our flow cytometry data and Sanger sequencing data, using 10-fold cross-validation, yielding a value of predicted mRFP production level for each data point. Dependent on the method used (LASSO or RFR) and the featurisation method, the prediction accuracy somewhat varies. However, all methods can predict protein levels reasonably well, with Pearson correlation coefficients ranging from 0.546 up to 0.776 (Figure 3, Table S2).

**Figure 3.**
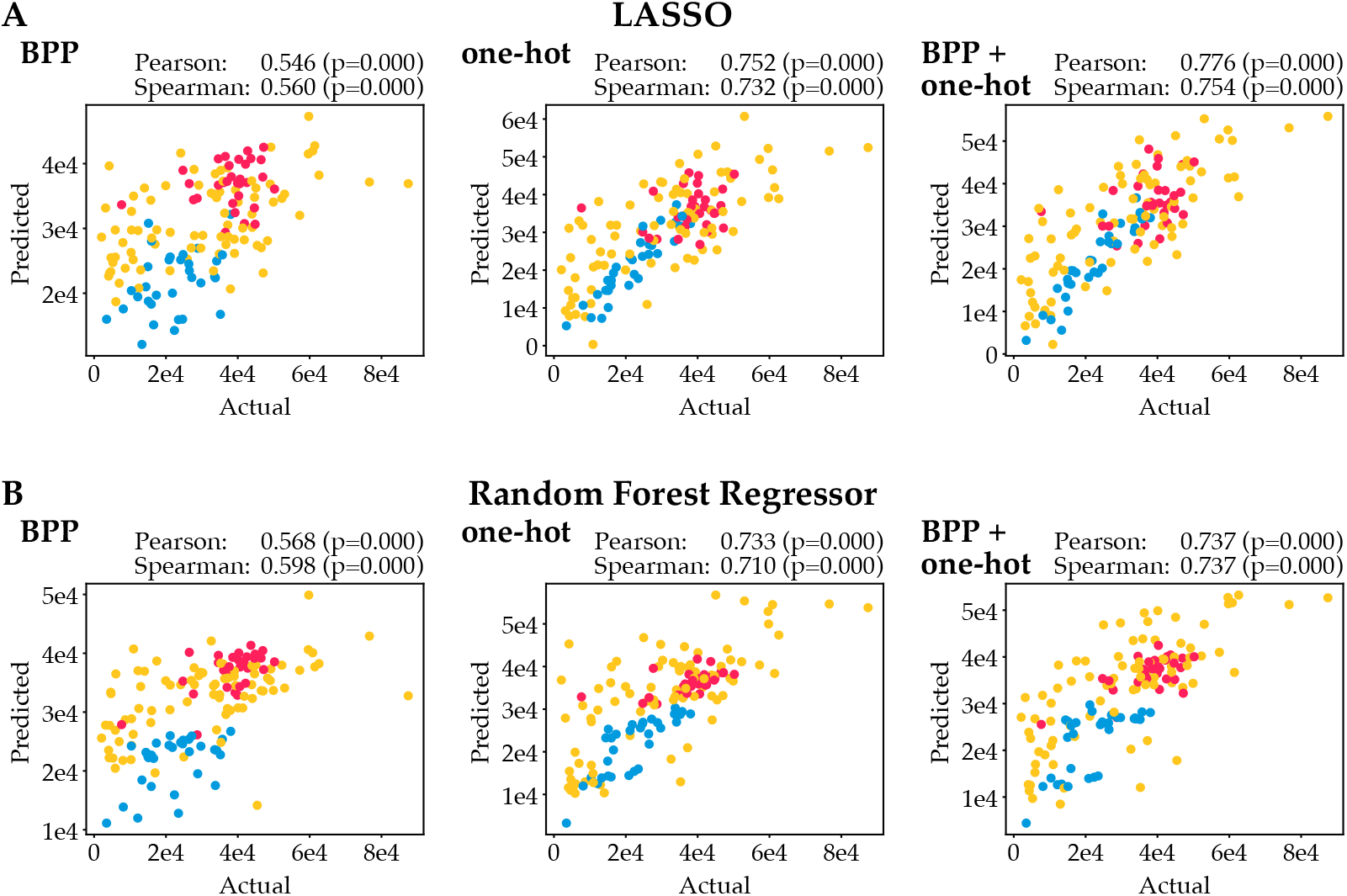
Actual expression data vs predicted expression using various machine learning algorithms and featurisations. Blue, yellow and red points indicate data points from the leave-out test sets for the CAI_L_, CAI_M_ and CAI_H_ libraries respectively. (**A**) Actual expression data vs predicted expression using LASSO. (**B**) Actual expression data vs predicted expression using the Random Forest Regressor (RFR). Regressor accuracies were evaluated through 10-fold cross-validation.

The predictive strength of the one-hot encoding featurisation is stronger than that of the BPP featurisation. This is not unexpected, as BPP featurisation assumes mRNA secondary structures are the only cause of expression variance, and base pairing featurisation is done based on an mRNA secondary structure model (ViennaRNA), which likely cannot perfectly predict exact base pairing. Still, BPP featurisation yields reasonable performance, which suggests that mRNA structures are a key factor in predicting protein production levels. One-hot encoding captures all information in the sequence and expectedly gives substantially better performance. Combining one-hot encoding with BPP featurisation does not substantially improve predictions, in line with our expectation that all information on base pairing probability should already be captured by one-hot encoding. No large differences in performance between the two regression methods LASSO and RFR are observed. When only BPP is used as feature, RFR performs slightly better than LASSO; when one-hot or one-hot + BPP are provided, LASSO slightly outperforms RFR.

### Bases surrounding the start codon and the RBS are most predictive of translation efficiency

Next, we assessed which features, and by extension which bases, are most predictive for translation efficiency. We did this by extracting the coefficients for LASSO and the feature importances for RFR and plotting them against sequence position (Figure 4). For the BPP featurisation information can be obtained for every nucleotide, including fixed nucleotides in the CDS and the UTRs, as they are still involved in the overall secondary structure formation. For the one-hot featurisation however, the information is only limited to changing nucleotides and can therefore be plotted per codon as only every third base changes.

**Figure 4.**
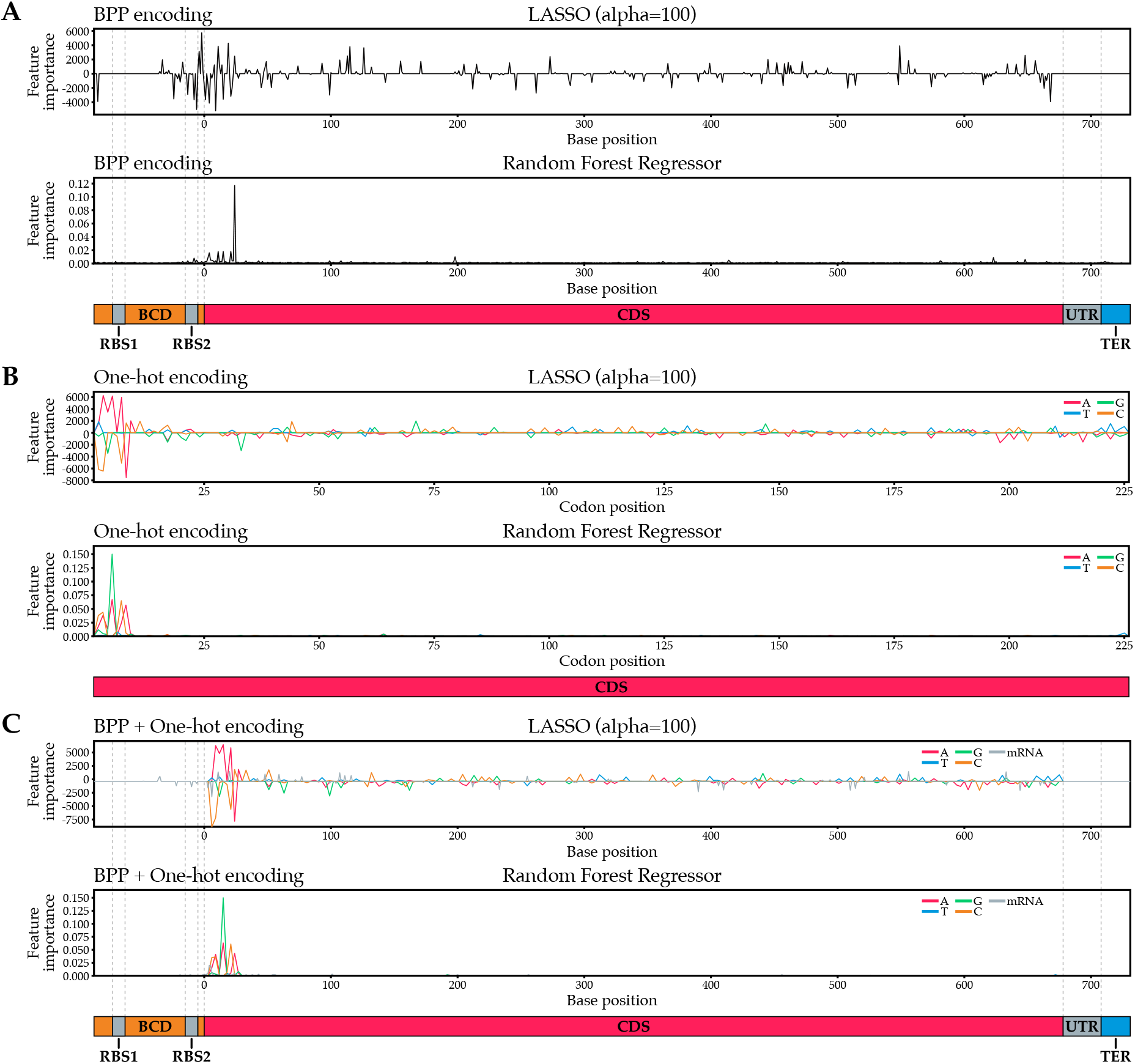
Feature importances for various machine learning algorithms and featurisations. LASSO feature importances are coefficients: a positive coefficient indicates a positive correlation between a base and translation efficiency, a negative coefficient indicates a negative correlation. In RFR, feature importances are always positive and therefore not indicative of the directionality of the correlation. (**A**) Feature importances for algorithms using BPP featurisation. (**B**) Feature importances for algorithms using one-hot encoding. Since only every third one-hot encoded base of the coding sequence varies, only every third base of the coding sequence was plotted. (**C**) Feature importances for algorithms using BPP + one-hot featurisation.

Overall, we found that independent of the algorithms used, the most predictive bases were always close to the start of the CDS, including the 5-10 bases before the start codon for BPP featurisations and the first 25 bases following the start codon for all featurisations. In comparison, the remaining codons play a minimal role in predicting translation efficiency. This strongly suggests that mRNA secondary structure or other factors around the start of the coding sequence play a dominant role in determining translation efficiency, in agreement with previous studies which found the same effect for GFP and attributed this to the necessity for an unobstructed RBS (2, 4). However, since we used a BCD system in the 5’ UTR, which should in theory reduce RBS obstructions prior to ribosome binding by straightening out the mRNA with a leading ribosome, the importance of the area surrounding the RBS was somewhat unexpected in our study. The BCD may be unable to completely resolve inhibitory secondary structure effects, or other factors at the start of the coding sequence may still play a dominant role.

Interestingly, the LASSO regressor using BPP for featurisation assigns positive coefficients to the bases immediately succeeding the RBS2 in the BCD region (Figure 4A, first panel). A positive coefficient indicates that involvement in mRNA secondary structures at this position is positively correlated with gene expression levels, which seems counterintuitive. In contrast, the bases in the RBS2 itself and the 5’ of the coding region are overwhelmingly assigned negative coefficients, which supports the hypothesis that minimal secondary structure surrounding the RBS is beneficial for high protein production levels. In concordance, the presence of A or T bases in the 5’ of the coding region, particularly ‘A’, is strongly positively correlated with protein production levels, while the presence of G or C bases tends to be negatively correlated with protein production levels in this region (Figure 4B, first panel). As A-T base pairs, and their A-U equivalents in mRNA, only form two hydrogen bonds versus three in G-C base pairs, the resulting secondary structures are weaker, and as a result, the RBS may be more accessible.

### A sequence window covering first 8 codons can predict expression well

To further substantiate our finding that bases surrounding the start codon dictate translation efficiency, we trained regressors on sliding windows of 10, 20, 30, or 40 bases to visualise which regions of the mRNA were most predictive of translation efficiency. For each sliding window, we performed a 10-fold cross-validation and plotted the correlation between actual expression data and the predicted expression data as a function of the position of the sliding window (Figure 5). Clear peaks of increased predictive power can be observed around the start codon, which corroborates our earlier finding that this region primarily dictates translation efficiency. This is especially apparent in models trained with one-hot encoded features and BPP + one-hot encoded features. Specifically, the 20 nucleotides surrounding base 15 (bases 6-25) are necessary for high prediction accuracy (Figure 5B, C, window size 20; Figure S5, S6). This window covers codons 2-8 and logically does not cover the start codon or the first two nucleotides of the second codon, as these are constant in our design and thus cannot have any predictive power in one-hot featurisation.

**Figure 5.**
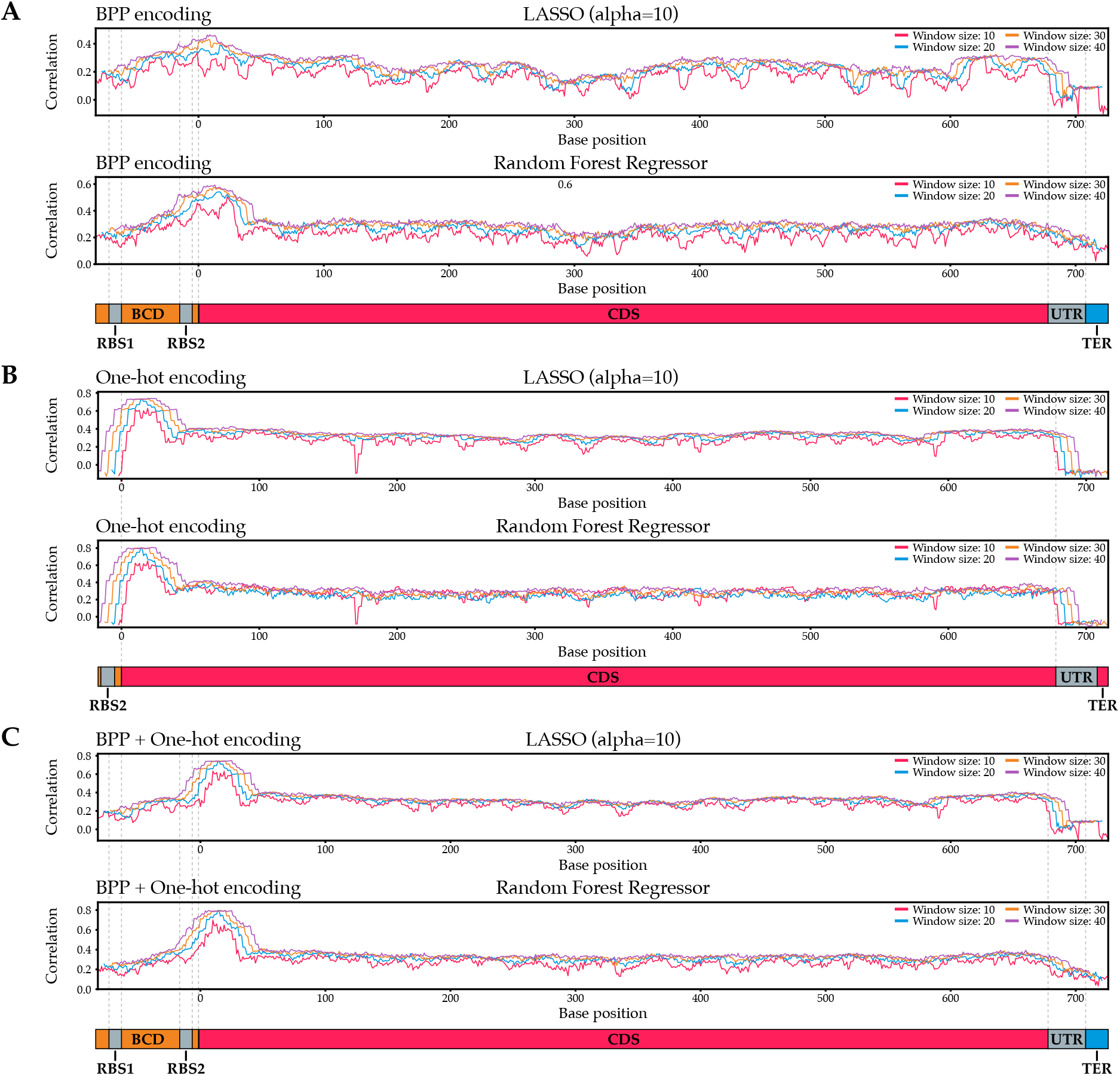
Predictive regions of translation efficiency in mRFP mRNA. The x-axis represents the central base of a sliding window of indicated lengths, the y-axis the correlation between actual expression data and the expression as predicted by a machine learning algorithm trained on solely that sliding window. (**A**) Predictive regions found by algorithms trained with BPP featurisation. (**B**) Predictive regions found by algorithms trained with one-hot encoding. As the one-hot encoded features for the UTRs are constant and thus contain no predictive information, windows that only contain residues in the UTR were omitted. (**C**) Predictive regions found by algorithms trained with BPP + one-hot featurisation.

It should be noted that the remainder of the CDS, while a lot less predictive than its 5’ region, still holds some predictive power (Pearson correlation ~ 0.4). We ascribe this to the inclusion of the CAI_H_ and CAI_L_ libraries in our dataset. These datasets use completely different sets of codons, meaning that the machine learning approaches can infer from small sequence windows which library a sequence originated from. Since data points from the CAI_H_ library on average display higher expression levels than data points from the CAI_L_ library, we attribute the non-zero Pearson correlations observed for windows downstream of the 5’ end of the CDS mostly to the algorithm’s ability to detect the library of origin of CAI_H_ and CAI_L_ data points. Inspection of scatter plots for windows in these regions confirmed this (Figure S7).

We also observed some ‘dips’ in predictability performance in the sliding window analysis. One such dip can be seen for small window sizes in the 3’ UTR with the BPP featurisation and LASSO regressor (Figure 5A, C). This region is very invariable both in terms of sequence and secondary structure: since the terminator almost always forms a strong secondary structure, the bases directly before it are less likely to be involved in secondary structures. As a result, the BPP features representing this region hold practically no information. The effect is exacerbated for small windows, as they are less likely to capture predictive residues upstream or downstream of an information-devoid region. In contrast, the secondary structure of the terminator itself does appear to be slightly informative. A perhaps unlikely but possible explanation could be that certain codon sequences interfere with the terminator stem formation and thereby influence mRNA stability. However, it is important to keep in mind that correlations between actual and predicted expression data for regressors trained on this region are still extremely low. Therefore, while the 3’ UTR region holds some information, it is not likely to be very influential.

A second dip is located around base 165 and 166 for regressors using one-hot-encoded featurisations (Figure 5B). This information valley is caused by an unusually constant region in the mRFP gene, particularly in the CAI_H_ library, due to the low codon variability of local amino acids and a fixed boundary region of two assembly blocks. This is an artefact of our method, and hence not a biologically relevant observation. This dip is not observed for featurisation methods that also include base pairing probabilities, as base pairing interactions of constant regions with other bases can still be informative.

To better understand which sequence elements in the 20-base window surrounding base 15 affect translation efficiency, we plotted feature importances for each regressor trained on this window (Figure S8). From this, we inferred that especially at position 15 (codon 5), low probabilities of involvement in mRNA secondary structure are predictive of high expression. This is in line with the current consensus that minimal mRNA secondary structure surrounding the 5’ end of the coding region is conducive to efficient translation. In the case of mRFP, this low base pairing probability seems to be primarily achieved by placing an ‘A’ at position 15 (Figure S8B, C).

Of all our regressors, six random forest regressors outperformed the rest (Figure S5, S6). As these six regressors were comparable in performance, we selected the model that used the fewest features, and retrained the model using our full training set. We then plotted actual expression data against predicted expression for each data point in our leave-out test set. This revealed a very strong correlation (Pearson r = 0.762) for all three libraries (Figure 6), which demonstrates that mRFP protein production can be correlated extremely well to sequence by just looking at bases 6-25 of the entire coding sequence.

**Figure 6.**
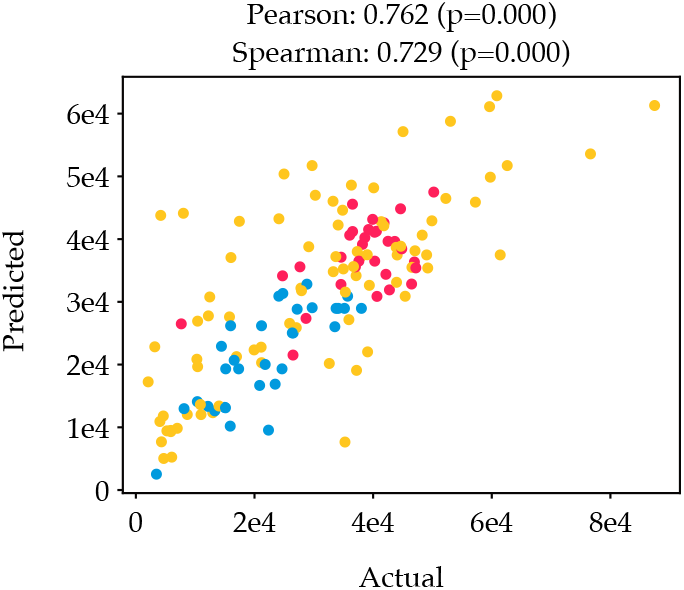
Actual vs predicted expression for one of our six best performing regressors. The RFR regressor on one-hot featurisation and a window size of 20 located at base 15 showed a high correlation between actual and predicted expression (Pearson correlation 0.762).

Altogether, our results show that while the CAI on average influences gene expression (Figure 2), the majority of the translation regulation arises from codon usage in the 5’ of the CDS. Because translation initiation is the major rate-limiting factor, the effects of codon usage throughout the gene will be less apparent. This is also exemplified by our finding that the highest expressing variants originate from our CAI_M_ library. This library contains more codon variance than the CAI_H_ library (Table S1) which is particularly important for the 5’ of the CDS. Due to the partial black box nature of machine learning, design rules for the 5’ CDS are not yet apparent. However, it is clear that if high protein production is desired the focus should be on the start of the coding sequence in combination with the 5’ UTR, in *E. coli* and probably in other bacterial hosts. Typically, codon optimisation algorithms and approaches optimise for parameters such as CAI over the full CDS, but ignore the 5’ UTR sequence and thereby possibly detrimental secondary structures. We suggest a shift in these approaches to specifically tackle optimisation of the first ~10 codons and develop experimental and *in silico* tools in this direction to make synthetic gene designs more productive.

## AVAILABILITY

MEW: the mRNA Expression Wizard is available at https://github.com/BTheDragonMaster/mew.

## ACKNOWLEDGEMENT

We acknowledge Markus Jeschek and Sjoerd Creutzburg for fruitful discussions and feedback on this work, as well as Rob Joosten and Christian Sudfeld for experimental support for flow cytometry and FACS operation.

