## Supplementary Figures and Tables for "Revealing determinants of translation efficiency via whole-gene codon randomisation and machine learning"

### SUPPLEMENTARY DATA

**Table S1. Theoretical and experimental sequence spaces for CAIL, CAIM and CAIH libraries.**

| Library | Theoretical sequence space | Experimental sequence space |
|---|---|---|
| $CAI_L$ | $4.47 \times 10^{49}$ | $6.22 \times 10^{38}$ |
| $CAI_M$ | $3.19 \times 10^{104}$ | $3.68 \times 10^{93}$ |
| $CAI_H$ | $2.01 \times 10^{53}$ | $4.50 \times 10^{48}$ |

**Table S2. Pearson and Spearman correlations for LASSO and RFR machine learning models trained on the full mRNA sequence using different featurisation methods**.

| Model | Featurisation | Pearson correlation (p-val) | Spearman correlation (p-val) |
|---|---|---|---|
| LASSO | BPP | 0.546 (0.000) | 0.560 (0.000) |
| | one-hot | 0.752 (0.000) | 0.732 (0.000) |
| | BPP + one-hot | 0.776 (0.000) | 0.754 (0.000) |
| RFR | BPP | 0.568 (0.000) | 0.598 (0.000) |
| | one-hot | 0.733 (0.000) | 0.710 (0.000) |
| | BPP + one-hot | 0.737 (0.000) | 0.737 (0.000) |

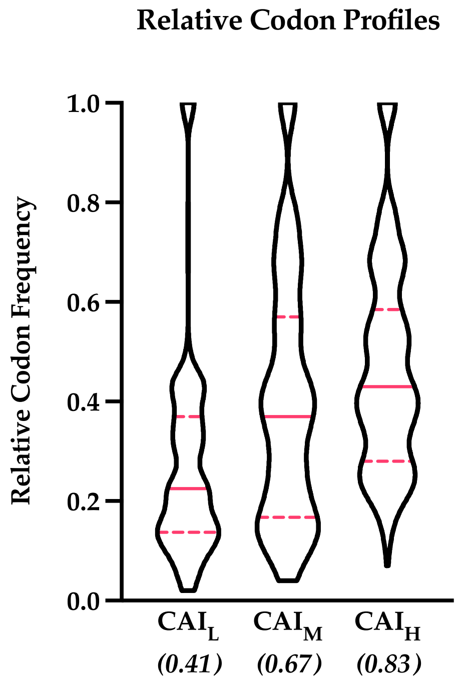


**Figure S1. Relative codon frequency profiles of the $CAI_L$, $CAI_M$ and $CAI_H$ libraries**. The solid red line indicates the median, dashed red lines indicate the quartiles. The CAI of each library is given in italics.

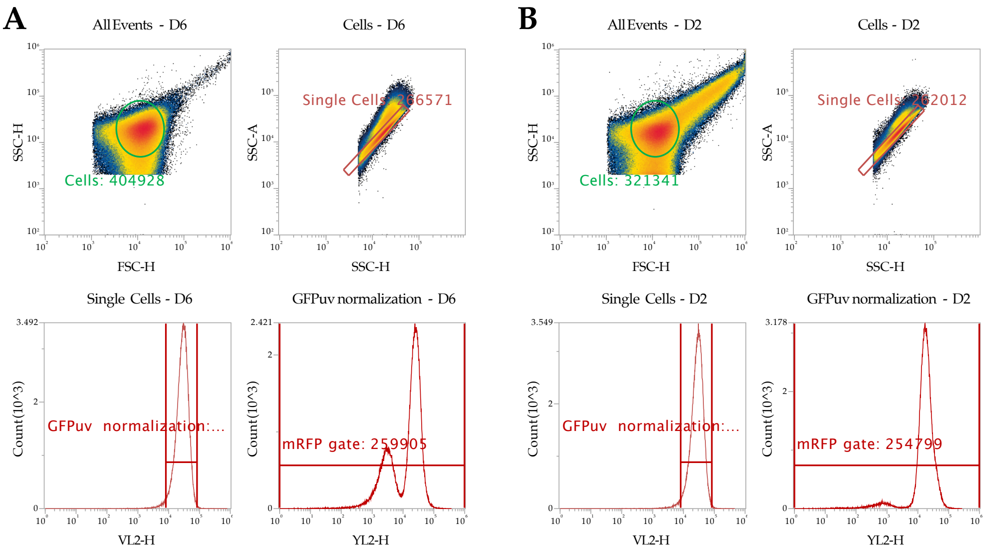


**Figure S2. Example of non-passed clones for double population and increased cell size/clumping**. (**A)** A flow cytometry environment where a clear double population is present in the mRFP expression (bottom right panel). (**B)** A flow cytometry environment where there is an increase in cell size or clumping of cells (top left panel, difference becomes apparent when compared to (**A**). The reason for this phenomenon is unknown.

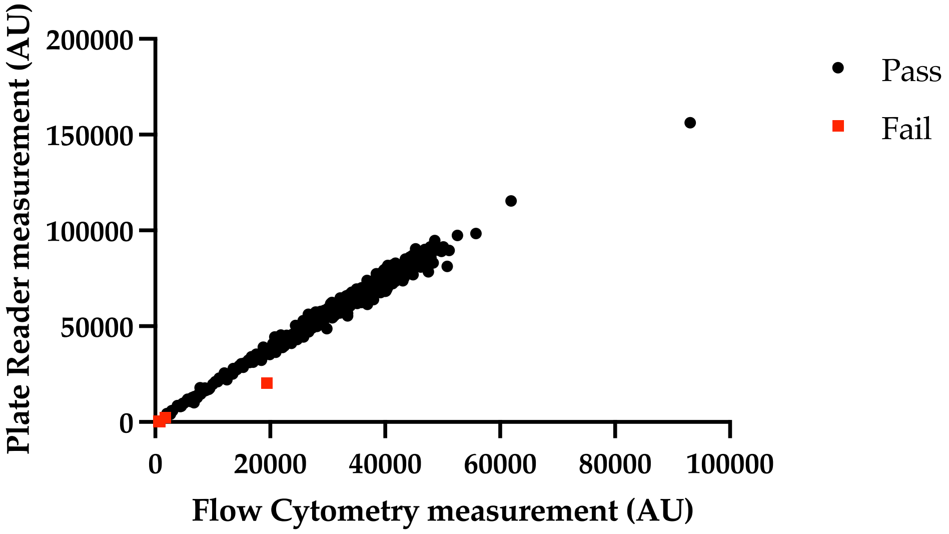


**Figure S3. Flow cytometry vs plate reader data**. Red points deviated more than 25% from the average ratio between all points and were excluded in the quality assurance check.

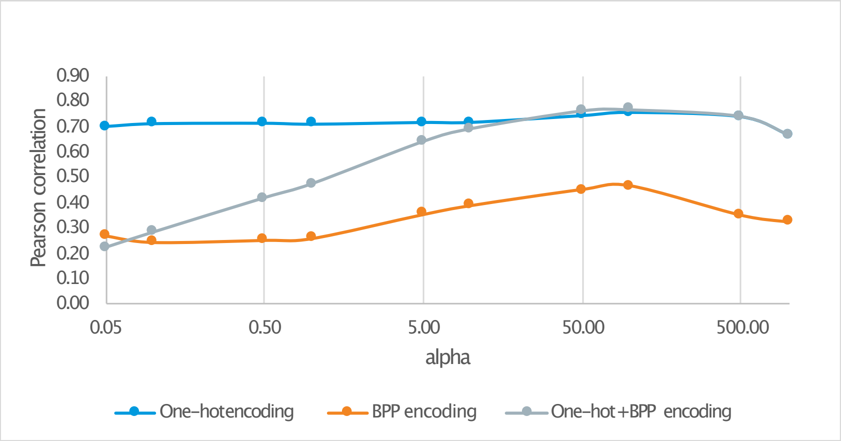


**Figure S4. Model performance with alpha for LASSO regressors trained on full-length sequences.** Model performance peaked at an alpha of ~100.0 for all featurisation methods

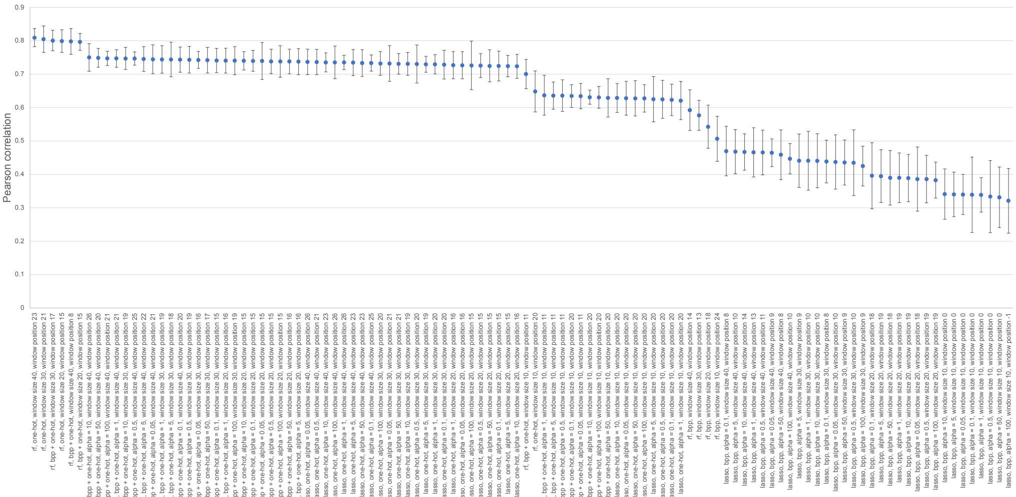


**Figure S5. Performance of machine learning models trained on best-performing sequence windows per parameter set**.

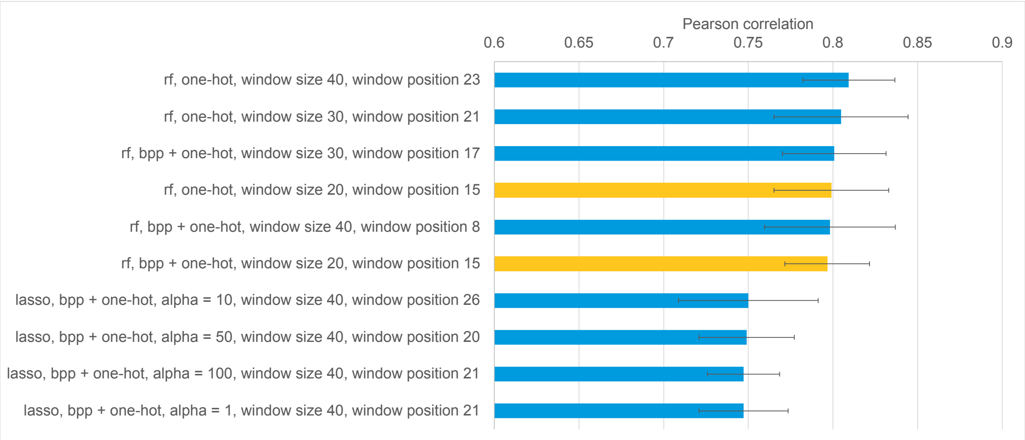


**Figure S6. Performance of machine learning models trained on best-performing sequence windows per parameter set, best 10 models.** Error bars show standard deviation of Pearson correlation across 10 cross-validation sets. The six best-performing models perform comparably.

Models highlighted in yellow use the narrowest sequence window (20 bases). As these models use fewer features without losing predictive power, this sequence window was considered the most important in determining protein production efficiency.

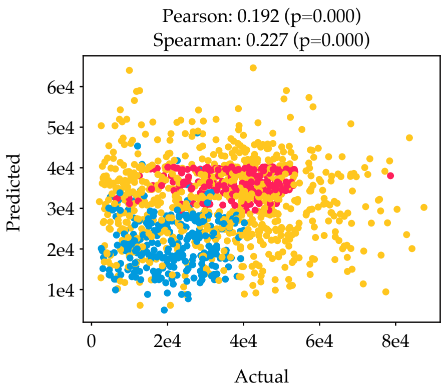


**Figure S7. Actual flow vs flow as predicted by a random forest regressor (one-hot encoded) through cross-validation on window 251 (window size 20).** The clear divide between the $CAI_L$ (blue) and $CAI_H$ (red) libraries shows that the model's (limited) predictive power relies on its ability to distinguish between these two libraries. Since these libraries use completely different codon sets, and since on average data points from the $CAI_H$ library display higher expression than data points from the $CAI_L$ library, it makes sense that the model is able to infer from just a small sequence window which library the sequence originated from. This accounts for the non-zero Pearson correlation we observe for most sequence windows.

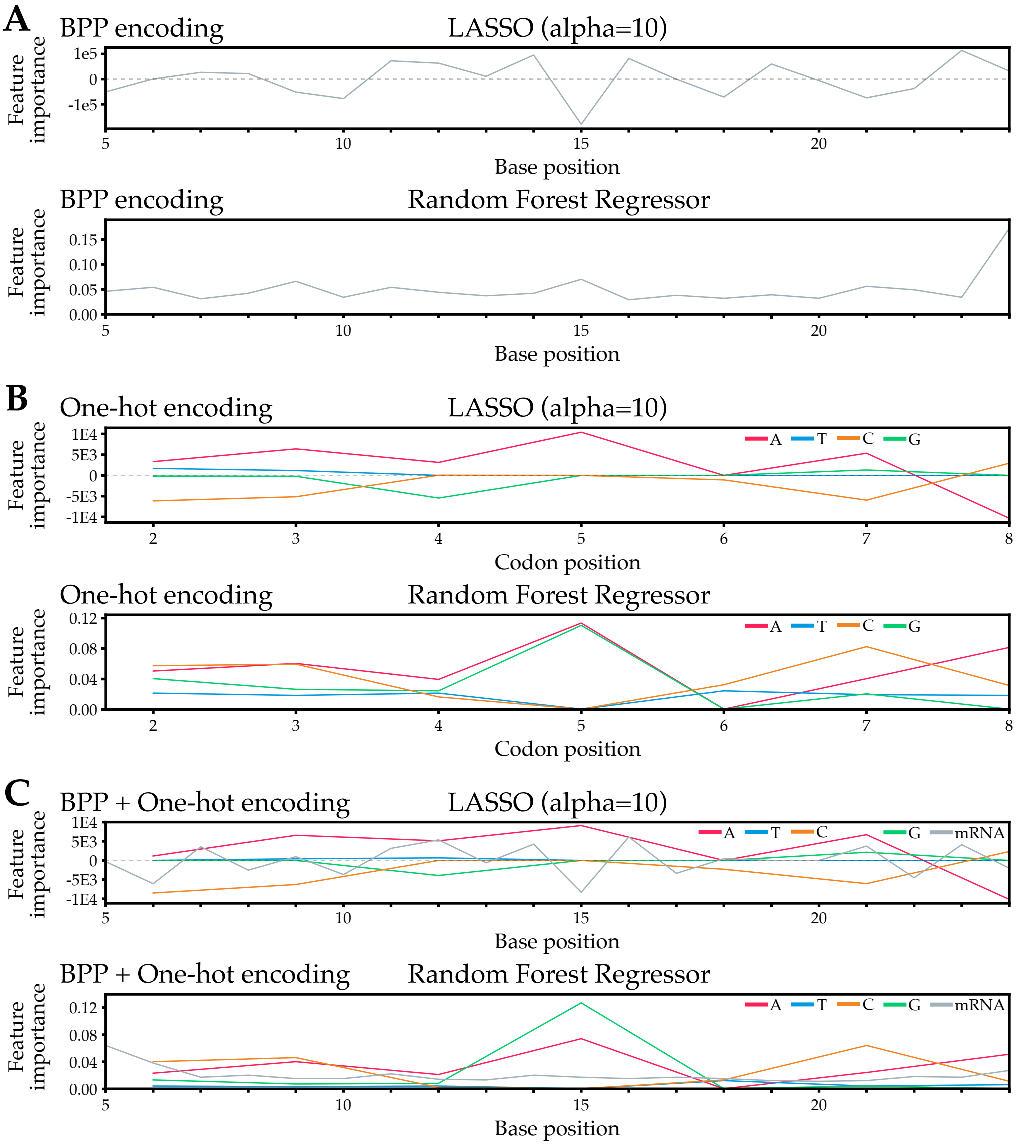


**Figure S8. Feature importances for various machine learning algorithms and featurisations trained on a 20-base window around base 15**. LASSO feature importances are coefficients: a positive coefficient indicates a positive correlation between a base and translation efficiency, a negative coefficient indicates a negative correlation. In RFR, feature importances are always positive and therefore it contains no information about the directionality of the correlation. (a) Feature importances for algorithms using BPP featurisation. (b) Feature importances for algorithms using one-hot encoding. Since only every third one-hot encoded base of the coding sequence varies, only every third base of the coding sequence was plotted. (c) Feature importances for algorithms using BPP + one-hot featurisation.
